# Metabolic insights into microbially induced calcite formation by Bacillaceae for application in bio-based construction materials

**DOI:** 10.1101/2024.11.12.623140

**Authors:** Michael Seidel, Charlotte Hamley-Bennett, Bianca J. Reeksting, Manpreet Bagga, Lukas Hellmann, Timothy D. Hoffmann, Christiane Kraemer, Irina Dana Ofiţeru, Kevin Paine, Susanne Gebhard

## Abstract

Microbially induced calcite precipitation (MICP) offers promising solutions for sustainable, low-cement infrastructure materials. While it is known how urea catabolism leads to biomineralization, the non-ureolytic pathways of MICP are less clear. This limits the use of the latter in biotechnology, despite its clear benefit of avoiding toxic ammonia release. To address this knowledge gap, the present study explored the interdependence between carbon source utilization and non-ureolytic MICP. We show that acetate can serve as the carbon source driving calcite formation in several environmental Bacillaceae isolates. This effect was particularly clear in a *Solibacillus silvestris* strain, which could precipitate almost all provided calcium when provided with a 2:1 acetate-to-calcium molar ratio, and we show that this process was independent of active cell growth. Genome sequencing and gene expression analyses revealed an apparent link between acetate catabolism and calcite precipitation in this species, suggesting MICP may be a calcium stress response.

Development of a simple genetic system for *S. silvestris* led to deletion of a proposed calcium binding protein, although this showed minimal effects on MICP. Taken together, this study provides insights into the physiological processes leading to non-ureolytic MICP, paving the way for targeted optimization of biomineralization for sustainable materials development.

## Introduction

The construction industry is, after transport, the second largest contributor to our current carbon emissions, with cement production alone contributing approximately 8% of anthropogenic CO2 release (Farfan et al. 2019). Alternative means of either replacing cement in construction materials or expanding the useful lifespan of cement-based infrastructure are therefore highly desirable from both an environmental and a sustainability viewpoint. The ability of many bacteria to precipitate calcium carbonate in the form of calcite has been gaining interest over recent years for its potential use in construction materials (Castro-Alonso et al. 2019; Justo-Reinoso et al. 2021; Seifan and Berenjian 2018). Calcite is the main component mineral of natural limestone, hence its production by bacteria may offer a green route to low-cement or cement-free materials.

One application of microbially induced calcite precipitation (MICP) that has seen considerable progress over the past decade is so-called “self-healing” concrete. While concrete is extremely strong under compression and very durable under ideal conditions, many environmental stresses or bending strains can lead to formation of microcracks. These cracks increase water permeability, leading to corrosion of steel reinforcements and reducing the service life of structures (Richardson et al. 2020; Seifan and Berenjian 2018). In self-healing concrete, bacteria are encapsulated, usually in spore form, and added directly to the concrete during casting, together with nutrients and often calcium salts (Alazhari et al. 2018; Seifan and Berenjian 2018; Richardson et al. 2020; Justo-Reinoso et al. 2021; Bagga et al. 2023). Once cracks occur, the encapsulating material of the bacteria breaks to release the cells, while ingress of water dissolves the nutrients and triggers spore germination.

Formation of calcite by the activated bacteria eventually seals the crack and restores water- tightness of the material to prevent further damage (Alazhari et al. 2018; Seifan and Berenjian 2018; Richardson et al. 2020; Justo-Reinoso et al. 2021).

Other potential applications of MICP include consolidation of soils, bioremediation of environmental heavy metal or radionuclide contaminations, or the development of Engineered Living Materials (Srubar 2021; Fu et al. 2023; Dikshit et al. 2022; Kumar et al. 2023). The diversity of technologies that are seeking to exploit the ability of bacteria to form water-insoluble minerals highlights the potential of MICP for the development of sustainable future materials. Despite the intensity of biotechnological research and development of MICP-based applications, we have a surprisingly limited understanding of the molecular mechanism that facilitate biomineralization. Such knowledge will, however, be required to select the best-suited bacterial species and optimise the applications for maximal mineral formation.

The majority of applications using MICP to date have been focussed on ureolytic bacteria. Here, the enzyme urease catalyses the hydrolysis of urea to ammonia and carbonic acid. The ammonia causes a strong increase in pH, which shifts the carbonate equilibrium from carbonic acid to carbonate ions. In the presence of Ca^2+^ ions, the availability of carbonate and the high pH values lead to precipitation of insoluble calcium carbonate (Hoffmann et al. 2021). Mineral precipitation resulting from ureolysis therefore has been described as a ‘passive’ process, based on induced chemical changes in the macroenvironment surrounding the bacteria (Castanier et al. 1999). While ureolysis leads to fast and strong biomineralization, the production of two moles of ammonia per mole of urea can lead to excessive nitrogen loading in the environment and toxic effects on animals, particularly in aquatic ecosystems (European Environment Agency 2019).

An alternative MICP mechanism that has seen promising developments, particularly in self-healing concrete, does not require the presence of urea and is usually simply described as “non-ureolytic”, reflecting our limited understanding of the drivers of biomineralization. While this form of MICP occurs more slowly than the ureolytic process, non-ureolytic bacteria have been used successfully in self-healing concrete (Jonkers et al. 2010; Justo-Reinoso et al. 2021; Bagga et al. 2023). Moreover, we could show that several non-ureolytic bacteria isolated from limestone environments were capable of the same high yields of MICP as their ureolytic counterparts (Reeksting et al. 2020). When tested for self-healing of cracked cement mortars, they gave a comparable, if not better, performance, showing their feasibility for industrial application (Reeksting et al. 2020).

Broadly, MICP in non-ureolytic bacteria is thought to rely on carbon dioxide evolved from bacterial metabolism of organic carbon compounds. In aqueous conditions, the resulting presence of dissolved inorganic carbon (DIC), i.e. the sum of carbonic acid, bicarbonate and carbonate ions, can cause the formation of calcium carbonate when high concentrations of calcium ions are provided (eq. 1-3) (Hoffmann et al. 2021; Schreiberová et al. 2019; Mondal and Ghosh 2019).

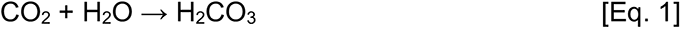

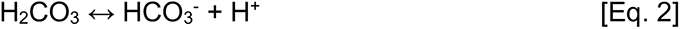

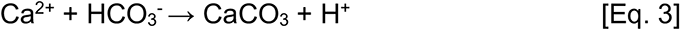

It has also been proposed that active maintenance of a physiological calcium balance in the bacteria could play a role (Hammes and Verstraete 2002). Intracellular calcium concentrations are estimated to be in the range of 0.1–0.3 µM, which is 1000-fold less than typically found outside the cell, especially in MICP conditions (Norris et al. 1996; Dominguez 2004). In such high calcium environments, passive influx will lead to an accumulation of intracellular calcium. This is potentially damaging to the bacteria, which likely counteract the influx by using active exporters, e.g. Ca^2+^-ATPases, to reduce intracellular calcium concentrations to physiological levels. It has been theorised that such active export could lead to a localised accumulation of calcium ions near the cell surface, where they could react with the DIC from the bacterium’s carbon metabolism to form calcium carbonate minerals (Hammes and Verstraete 2002).

Taking these contributing components of bacterial metabolism together shows that non-ureolytic MICP is a much more complex process than ureolytic MICP. To understand how biomineralization occurs and design effective applications it is essential to gain detailed mechanistic insights. This involves studying the metabolic properties of non-ureolytic MICP bacteria. Additionally, it is important to understand how carbon and calcium metabolism are linked to mineral formation, both on a physiological and genetic level.

In this study, we set out to address this question by investigating the links between carbon metabolism and MICP in a subset of our previously isolated environmental, non-ureolytic MICP strains (Reeksting et al. 2020). Development of a defined medium based on acetate as the main carbon source allowed us to show that, in some species, carbon catabolism indeed was directly correlated with calcium carbonate formation. Whole genome sequencing of five isolates was used to identify candidate genes that may be involved in a physiological link between carbon metabolism, calcium detoxification and MICP. For the most MICP-active isolate, *Solibacillus silvestris* CGN12, we could show that calcium addition caused increased expression of genes for both acetate metabolism and calcium detoxification. To investigate this further, an initial system for the genetic modification of *S. silvestris* CGN12 was developed to delete a putative calcium binding protein and assess its impact on MICP. Overall, our data provide insights into the mechanisms of non-ureolytic MICP and experimental evidence that carbon catabolism and potentially calcium detoxification directly influence biomineralization.

## Results

### Calcium carbonate precipitation using acetate as a carbon source

From a metabolic perspective, ureolytic MICP is straightforward to understand, being largely linked to the hydrolysis of urea to carbonate and ammonia through the activity of the enzyme urease (Stocks-Fischer et al. 1999). For non-ureolytic MICP, the metabolic mechanisms leading to biomineralization are far less clear. Past studies have mainly used yeast extract-based media for assessing the ability for MICP and quantity of calcite minerals formed during growth of non-ureolytic bacteria, e.g. the model MICP species *Alkalihalobacillus pseudofirmus* (formerly *Bacillus pseudofirmus*) and *Sutcliffiella cohnii* (formerly *Bacillus cohnii)* (Justo-Reinoso et al. 2021). While growth in such complex media leads to reliable production of biomass and, in the presence of calcium ions, calcite formation, it does not provide insights into specific metabolic pathways used by the bacteria, e.g. linking carbon metabolism to calcite formation.

To address this, we set out to develop a more defined growth medium that would allow meaningful comparisons between performance of model species and uncharacterised environmental strains during growth on a specific carbon source. We focussed on the non-ureolytic environmental species from a previously assembled strain collection, most of which belonged to the genus *Bacillus* or close relatives (Reeksting et al. 2020). Furthermore, we based the growth medium on the composition of CSE medium, a chemically defined medium commonly used to grow *B. subtilis* (Commichau et al. 2008), but with modifications as detailed below.

Metabolic predictions based on species-level identifications by 16S rDNA sequencing obtained in our previous study (Reeksting et al. 2020) suggested that most of the strains would likely display some form of amino acid auxotrophy, so we decided to include a small amount of yeast extract (0.2%) to provide a source of not only nitrogen but also essential amino acids, vitamins, and trace elements. The provision of some yeast extract also counteracted the potential draw-back of a fully defined medium composed of many individual ingredients, which could make it impractical for industrial-scale applications in the future. For a suitable carbon source for non-ureolytic MICP, we considered that lactate or acetate are already commonly used in applications and therefore should be chemically compatible with cement-based technologies (Justo-Reinoso et al. 2021). Genomic comparison of pathways involved in carbon metabolism suggested that several isolated species likely lacked genes associated with lactate metabolism, discounting this as a potential carbon source. Acetate was therefore selected as the main carbon source for the medium, which we termed YA (for yeast extract and acetate), or YAC (yeast extract, acetate, calcium) medium. For routine assays, sodium acetate was provided at a concentration of 100 mM, with additional provision of 100 mM calcium nitrate where MICP was to be tested.

In preliminary tests, this medium was able to support robust growth of many of the initially selected isolates. Of these, we chose ten for further analysis (Table 1). These bacteria all showed mesophilic growth at 30°C. For some applications of MICP outside the laboratory it is, however, likely that bacteria with a lower temperature growth optimum are required. To this end, we repeated the isolation procedure described previously (Reeksting et al. 2020), but incubated the cultures at 7.5°C. One of the isolates obtained from this (strain Psy5), which could grow well in YA medium and showed robust psychrotrophic growth between 7.5 and 20°C, was selected for further analysis and included in our list of study species (Table 1). Finally, we included the two model species for non-ureolytic MICP, *A. pseudofirmus* and *S. cohnii*, for comparison and to provide a performance baseline.

**Table 1.** Overview of environmental isolates used for MICP analysis.

| Strain | Species<sup>a</sup> | Temperature preference | Reference |
| --- | --- | --- | --- |
| BA32 | *Metabacillus dongyingensis* | mesophilic | (Reeksting et al 2020) |
| BA41 | *Brevibacillus choshinensis* |  |  |
| BA42 | *Brevibacillus frigoritolerans* |  |  |
| BA43 | *Bacillus idriensis* |  |  |
| CG7-2 | *Lysinibacillus varians* |  |  |
| CGN12 | *Solibacillus silvestris* |  |  |
| CGN14 | *Jeotgalibacillus sp.* |  |  |
| PD1-1 | *Bacillus licheniformis* |  |  |
| UBN2 | *Bacillus frigoritolerans* |  |  |
| UBN3 | *Solibacillus isronensis* |  |  |
| Psy5 | *Peribacillus simplex* | psychrotrophic | This study |

<sup>a</sup> identified via 16S rDNA sequencing or available full genome sequences

For this selection of 13 strains, we assessed which species were capable of MICP using 100 mM acetate as a carbon source. The mesophilic isolates as well as *A. pseudofirmus* and *S. cohnii* were grown at 30°C for 6 days, while the psychrotrophic bacterium Psy5 was grown at 10°C for 12 days due to its slower growth rate. After this time, the amount of precipitated calcium carbonate was quantified. This showed that all tested species could precipitate calcium carbonate under the chosen growth conditions (Fig. 1A). The amount of calcium carbonate formed ranged from 20 mM in isolate CGN14 to 60 mM in isolate Psy5, with the two model species falling approximately in the middle of this range. Comparing these values to the theoretical maximal formation of calcium carbonate (100 mM), based on the amount of calcium salt provided in the growth medium, showed that only about half of the calcium became biomineralized in these experiments. This suggested that MICP activity was limited by an unknown factor under the chosen conditions.

**Figure 1.**
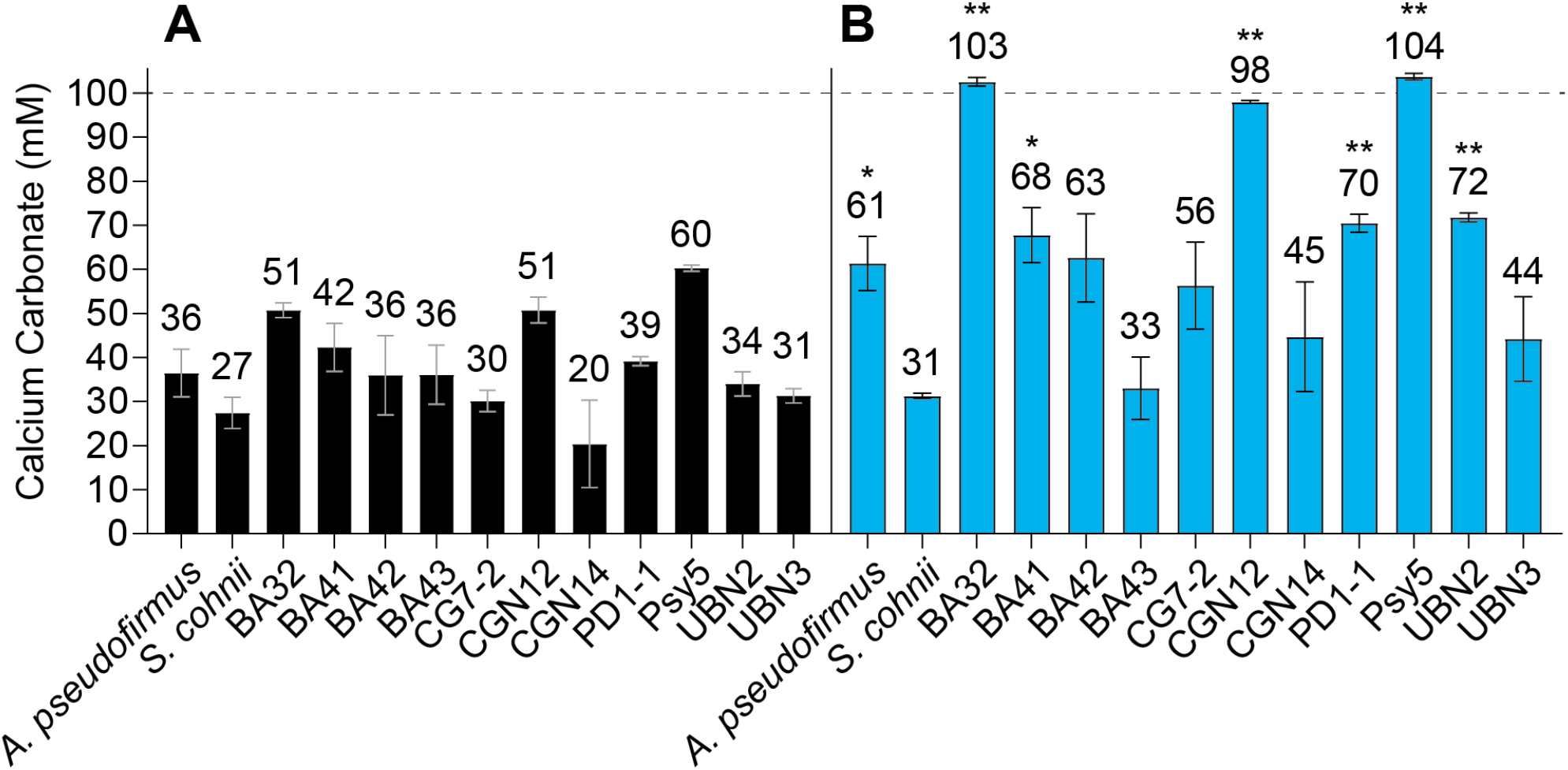
Comparison of calcium carbonate precipitation in YAC medium. *A. pseudofirmus, S. cohnii* and 11 environmental isolates were grown in YAC medium containing 100mM calcium nitrate and either (A) 100mM or (B) 200mM sodium acetate. All cultures were grown for 6 days at 30°C except Psy5, which was grown at 10°C for 12 days, at which time the calcium carbonate precipitate was harvested and quantified. Numbers above the columns show the mean of calcium carbonate in mM harvested from 2-3 biological repeats. Error bars are the standard deviation of the mean. Results ofstatistical comparison ofdata in panel B to data in panel A for each strain by Student’s t-test are shown as * for *P*<0.05 and ** for *P*<0.005. The horizonal line visualizes the theoretical maximum of calcium carbonate precipitation based on 100mM available calcium nitrate. Results are shown as the mean and standard deviation from three biological repeats.

A likely candidate for the limiting factor was the provision of the carbon source, because metabolic activity under aerobic conditions should convert acetate to CO2, resulting in an increased concentration of DIC, one of the requirements for calcium carbonate formation (Talaiekhozan et al. 2013). To test if this was the case, the concentration of acetate was doubled, and the experiment repeated under otherwise identical conditions (Fig. 1B Upon doubling the acetate concentration in the medium, the environmental strains BA32, CGN12, and Psy5, showed a strong and significant increase in calcium carbonate production, precipitating all of the available calcium. The strains PD1-1 and UBN2 also showed a moderately increase in calcium carbonate production, while the remaining strains did not show a significant increase.

Apart from demonstrating that the MICP efficiency in the initial experiment for some strains indeed had been limited by the provision of the carbon source, these results showed that increasing the acetate concentration had different effects across the bacteria tested. In CGN12, BA32 and Psy5, doubling the acetate concentration doubled the MICP yield, suggesting a directly proportional relationship between carbon source and biomineralization. At the other extreme, MICP by BA43 and *S. cohnii* was not affected by changing the acetate concentration. In these species, it appears that MICP is not immediately linked to the breakdown of this carbon source. Elucidating the exact reasons for this was beyond the scope of this study, but it is plausible that the precipitation activity of each species might be improved by optimising the provision of nutrients depending on specific needs.

### Acetate metabolism as a means of controlling calcite precipitation

To further explore the relationship between acetate concentration and calcium carbonate precipitation, we decided to focus on the three strongest precipitators, CGN12, BA32 and Psy5, as well as the better-understood model species *A. pseudofirmus* and *S. cohnii*. HPLC-analysis of culture supernatants from day 6 post-inoculation (12 days for Psy5), as in the experiments shown in Fig. 1, had not shown any detectable acetate remaining, indicating the carbon source had been consumed completely at some point during the incubation time.

Hence, we reduced the growth time to 3 days (6 days for Psy5) so that the rate of acetate utilisation could be determined.

Precipitation profiles were produced at different calcium nitrate concentrations (either 50 mM or 100 mM) while keeping the acetate concentration fixed at 100mM. As a general trend, changing only the calcium nitrate concentration appeared to have little effect on the precipitation of calcium carbonate (compare black hatched to black solid bars in Fig. 2A). To test if acetate was again the limiting factor, we doubled the amount of acetate (200 mM; Fig. 2A, blue bars). Increasing the acetate concentration in the presence of 50 mM calcium salt had no effect (compare blue hatched to black hatched bars), showing that here MICP was limited by the provision of calcium ions. For *A. pseudofirmus*, *S. cohnii* and Psy5 grown in the presence of 100 mM calcium nitrate, there were no obvious changes in precipitation patterns compared to the lower acetate concentration (Fig. 2A, blue vs black bars). However, for BA32 and CGN12, doubling the acetate concentration led to increased precipitation at higher calcium concentration. This effect was most pronounced for CGN12, where the amount of precipiated calcium carbonate almost doubled with the increase in acetate (Fig. 2A, blue hatched vs blue solid bars). These data confirmed that there was a direct connection between acetate provision, calcium concentration and the amount of calcium carbonate biomineralization in CGN12 and BA32.

**Figure 2.**
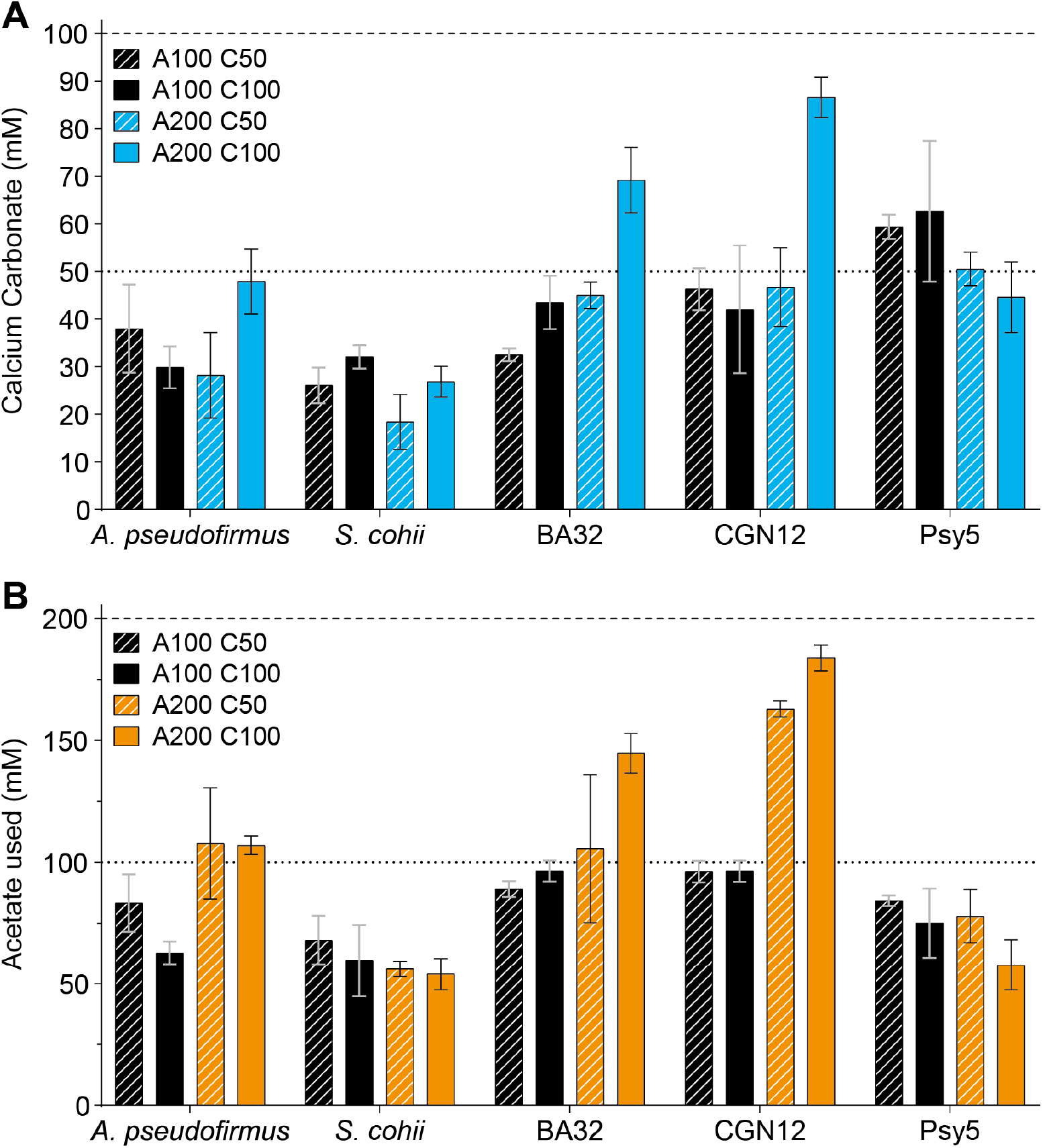
Comparison of calcium carbonate precipitation and acetate consumption between strains. Strains were grown in YAC medium with varied concentrations of acetate and calcium for 3 days at 30°C for all strains except Psy5, which was grown for 6 days at 10°C. A100 = 100mM, A200 = 200mM sodium acetate; C50 = 50mM, C100 = 100mM calcium nitrate. A. Quantification of calcium carbonate precipitation at the end of the experiment. Horizonal lines indicate the theoretical maximal calcium carbonate precipitation at 50mM (thin dashed line) and 100mM (thick dashed line) calcium nitrate. B. Acetate consumption (in mM) by the end of the experiment determined via HPLC analysis. Horizonal lines indicate the theoretical maximum amount of acetate utilisation at 100 mM (thin dashed line) or 200mM (thick dashed line) initial sodium acetate provision. Results are shown as the mean and standard deviation from three biological repeats.

To directly link the increases in precipitation with acetate consumption, acetate usage by each species under the chosen conditions was determined (Fig. 2B). *S. cohnii* showed similar levels of acetate utilization at 100 mM and 200 mM, not increasing usage in response to higher acetate levels, and also calcium concentration had no impact on acetate consumption. Actetate utilization in *A. pseudofirmus* increased slightly, but not proportionally, when more acetate was available (Fig. 2B, orange vs black bars). At lower acetate concentrations, calcium showed a slight inhibitory effect on acetate consumption by this bacterium (83 vs. 63 mM used), however this effect was not present at the higher acetate concentration (107 vs. 108 mM used). When provided with 100 mM acetate, CGN12 consumed almost all of the available acetate (96 mM used), which was unaffected by different calcium concentrations. By doubling the available acetate, acetate usage was nearly doubled (163 mM used), showing that this species’ metabolism was exclusively limited by carbon source provision in the YA medium. Increasing the calcium concentration led to a further small increase in acetate consumption by CGN12 (184 mM used). BA32 showed a similar pattern of usage to CGN12, degrading almost all of the available acetate (89-96 mM used) when 100 mM was provided. Similar to CGN12, at the higher acetate concentration, doubling the calcium concentration also stimulated acetate consumption by BA32 (106 mM vs 145 mM used).

These data supported our earlier observation that calcium precipitation by BA32 and CGN12 was directly driven by acetate consumption, while in the other species further aspects must be involved that are not easily explained with C- and Ca^2+^-source availability. Particularly in CGN12, it appeared that optimal biomineralization required a two-fold molar excess of acetate over calcium. A possible explanation for this seemingly fixed ratio could be the provision of DIC from acetate metabolism, which can then react with available calcium ions to form calcium carbonate minerals. This point is discussed further below.

Interestingly, Psy5, which had shown a clear acetate-dependency of biomineralization when tested over the original 12-day period (Fig. 1), did not demonstrate such an effect over the shorter incubation period used here. When provided with 100 mM calcium salt, it only produced about half the amount of theoretically possible mineral, regardless of how much acetate was provided (Fig. 2A). Consistent with this, quantification of acetate consumption also showed no change between the four growth conditions (Fig. 2B). It is likely that the lower incubation temperature and much slower growth of this species impacted on its metabolism and biomineralization in a way that halving the incubation period did not allow the completion of either acetate consumption or calcium carbonate formation.

To further investigate the links between acetate metabolism and calcium carbonate biomineralization, we focused on the environmental isolate CGN12 and the model species *A. pseudofirmus* for the rest of this study. The rationale for this selection was that CGN12 showed the strongest link between acetate consumption and MICP, while *A. pseudofirmus* at least showed a small increase in biomineralization at the elevated acetate concentration.

### Differences in precipitation kinetics between species

To gain a better understanding of the link between acetate metabolism and biomineralization on a temporal scale, we next determined the precipitation kinetics for CGN12 and *A. pseudofimus*. Both species were inoculated into YAC medium containing 100 mM calcium nitrate and either 100 mM or 200 mM sodium acetate, and cell growth as colony forming units ml^-1^ (CFU/ml), remaining acetate concentration and amount of precipitated calcium carbonate were determined after 1, 2, 3 or 6 days of incubation (Figure 3).

**Figure 3.**
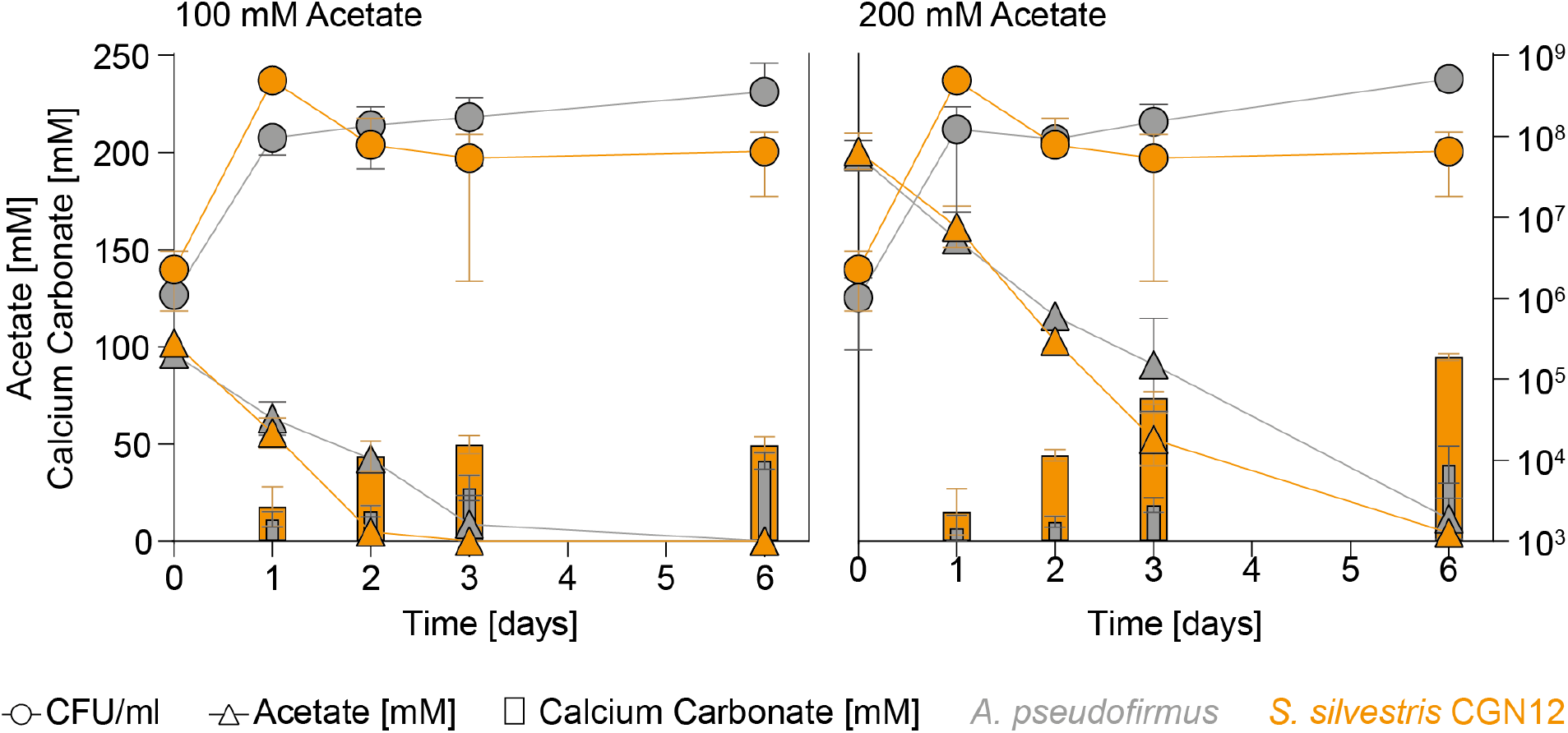
Precipitation kinetics of *A. pseudofirmus* and CGN12. Viable cell counts (CFU/ml), calcium carbonate precipitation and acetate consumption for *A. pseudofirmus* and CGN12 grown in YAC medium at different concentrations of sodium acetate were monitored over 6 days at 30°C with shaking (140rpm). The concentration of calcium nitrate was fixed at 100mM. Data are shown as mean and standard deviation of the mean for three biological replicates.

In agreement with our previous observations, CGN12 precipitated calcium carbonate depending on the available acetate, reaching maximal levels when all acetate was consumed (Fig. 3, orange symbols). This was after 2 days in the cultures with 100 mM acetate, after which precipitate amounts no longer increased. At the higher acetate concentration, the substrate was only fully consumed by day 6, and accordingly more mineral precipitate was formed until that time point. For *A. pseudofirmus*, acetate consumption was generally slower than for CGN12, requiring three days to consume the 100 mM concentration, and residual amounts still detectable after six days when 200 mM were initially provided (Fig. 3, grey symbols). As with CGN12, mineral production reached the highest levels at the time point where acetate was completely consumed, although overall lower levels of biomineralization were observed (approx. 40% of the theoretical maximum) with no increase observed at higher starting acetate concentration, consistent with the earlier assays.

Because acetate serves as a general carbon source in the medium, we needed to ensure that the apparent beneficial effect of acetate on calcium precipitation by CGN12 was not simply due to a general improvement of growth when more carbon source was available. To this end, we determined the Monod constant for CGN12 growth on acetate, i.e. the concentration that supported the half-maximal exponential growth rate, akin to Michaelis-Menten kinetics of enzymes (Pirt 1975). This resulted in values of KS = 69 mM (Monod- constant) and µMAX = 1.04 h^-1^ (maximal theoretical growth rate) (Figure S1), showing that both 100 mM and 200 mM acetate were considerably higher than the Monod constant, and only a marginal increase in growth should be expected from this change in concentration.

Interestingly, in both species cell growth (detected as viable cell counts) plateaued already after day 1 of incubation, at a time where acetate was still available and still being consumed. Cell growth was therefore limited by something other than the carbon-source, while biomineralization in CGN12 appeared only to be determined by the availability of sufficient acetate. Consistent with this observation, in both species, biomineralization was minimal after one day and continued to increase with longer incubation. This showed that mineral formation was linked to acetate catabolic activity, but not active cell growth, which is addressed in more detail next.

### Acetate drives MICP in stationary phase cells of *S. silvestris* CGN12

The observation that the majority of calcium carbonate formation occurred during the stationary growth phase of CGN12 next prompted us to investigate biomineralization specifically in this growth phase. This setup would allow us to track only acetate consumption and mineral formation, while cell counts and therefore biomass were held approximately constant. To this end, we inoculated five parallel small flask cultures with CGN12, each containing 50 ml of YA medium (180 mM sodium acetate), and allowed the cultures to reach stationary phase after 24 h of incubation. At this point, 100 mM calcium nitrate was added to each flask and incubation continued for another 24 h. For each time point, one entire culture was harvested to quantify precipitated calcium carbonate, viable cell counts (CFU/ml) and the remaining soluble calcium and acetate concentrations (Fig. 4).

**Figure 4.**
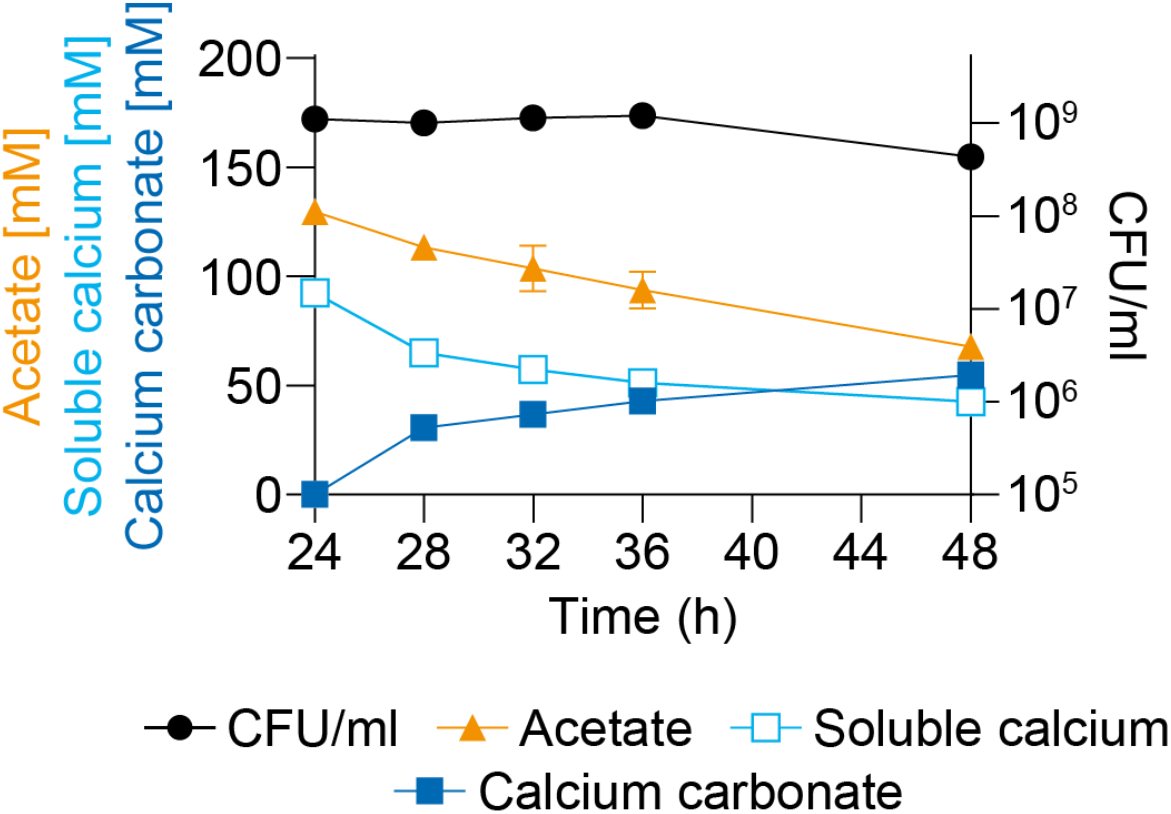
Acetate-driven biomineralization in stationary phase cells of *S. silvestris* CGN12. Cells were grown in five parallel flasks in YA medium with 180 mM acetate for 24 h to reach stationary phase. At this point, 100mM calcium nitrate was added aseptically, and the cultures were incubated for a further 24 h. At 4,8, 12 and 24 h after the addition of calcium nitrate, one entire flasks was harvested and precipitated calcium carbonate, viable cells (CFU/ml), and the concentrations of soluble calcium and acetate were quantified. Results are shown as mean and standard deviation from two biological repeats.

As seen before in the precipitation kinetics tests, at entry into stationary phase (24 h time point), the majority of the provided acetate remained, and acetate consumption proceeded at a steady rate over the following 24 h, while cell counts remained stable (Fig. 4, orange and black symbols). Interestingly, the concentration of soluble calcium showed a similarly steady decrease over the incubation period as for acetate. At the same time, we observed a gradual increase in insoluble CaCO3 mineral (Fig. 4, blue open and closed symbols). The appearance of the mineral accurately accounted for the amount of soluble calcium that had disappeared from the medium, with between 92% and 97% of total calcium recovered at each time point. As a negative control, an uninoculated control flask was also tested for precipitation after addition of calcium nitrate. The slightly alkaline pH of the medium resulted in the production of 1.6 mM (±0.1) calcium carbonate over the same 24 h period, which was negligible compared to the biological formation of mineral in the bacterial cultures.

The consistent rate of reduction in both acetate and soluble calcium concentrations during the time-course of this experiment suggested that the bulk of the provided acetate was needed during stationary phase and that at least some of the carbon from acetate degradation likely went towards calcium carbonate production via the formation of DIC.

### The yeast extract component of YAC medium does not significantly contribute to biomineralization

So far, we had assumed that the yeast extract in the growth medium did not significantly contribute to DIC provision. To confirm this, the individual contributions of acetate or yeast extract to precipitation of calcium carbonate were assessed in resting cell assays with CGN12. This format was necessary, because CGN12 was unable to grow in the defined medium when yeast extract was omitted, but it could maintain viability if cells pre-grown in YA medium were resuspended to high densities (OD600 = 4) in a solution that corresponded to YAC medium (50 mM calcium nitrate) but missing the yeast extract component. Following incubation for three days, cell viability was determined as CFU/ml, acetate consumption quantified by HPLC, and the precipitate harvested and quantified (Table 2).

**Table 2.** Resting cell precipitation assay of CGN12 under different concentrations of acetate.

| Acetate (mM)<sup>a</sup> | Average cell count at end of assay (CFU/ml)<sup>c</sup> | CaCO<sub>3</sub> produced (mM) | Acetate utilised (mM) | Stoichiometry (CaCO<sub>3</sub>: Acetate) |
| --- | --- | --- | --- | --- |
| 0<sup>b</sup> | 4.50E+08 | 0.0 (±0.0) | 0.0 (±0.0) | - |
| 50 | 1.30E+08 | 23.8 (±1.5) | 49.1 (±3.0) | 0.48 (±0.01) |
| 100 | 3.00E+07 | 47.5 (±0.5) | 102.4 (±5.1) | 0.47 (±0.02) |

<sup>a</sup> Medium composition corresponded to YAC medium with 50 mM calcium nitrate, but without the yeast extract component.
<sup>b</sup> YA base medium with 0.2% yeast extract provided as the sole carbon source.
<sup>c</sup> Cells were resuspended to a starting OD<sub>600</sub> = 4, corresponding to viable cell counts of approximately 5x 10<sup>8</sup> to 2x 10<sup>9</sup> cfu/ml on day 0. The end-point of the assay was day 3.

When the cells were provided with calcium nitrate and yeast extract but no acetate, no precipitation was observed after three days, demonstrating that neither the small amount of yeast extract present in the YAC medium nor carbon dioxide from the atmosphere were contributing relevant amounts of DIC for calcium carbonate precipitation. Hence, under the conditions tested, CGN12 appears to only use the carbon from acetate metabolism for calcium carbonate precipitation.

Where acetate was present, CGN12 was able to consume the entire provided carbon source during the three days of incubation. This acetate consumption did not result in further cell growth, because viable counts were slightly lower than where no acetate was provided (Table 2). Instead, acetate again appeared to be the limiting factor for precipitation of the provided calcium nitrate, closely recapitulating the earlier results from cells actively grown in YAC medium. As before, a two-fold molar excess of acetate over calcium was required for complete precipitation, because at 50 mM acetate, only approximately half the calcium was recovered as CaCO3, whereas it was nearly completely precipitated at 100 mM acetate (Table 2). Further comparison of CaCO3 production to acetate consumption showed that under both conditions the ratio of carbonate-to-acetate was almost identical at 0.48 (Table 2). Given that acetate is a C2-compound, this implies that approximately 25% of the carbon from acetate consistently went towards mineral formation, regardless of the initial amount of acetate provided.

### Acetate as the main carbon source in self-healing concrete

As MICP in CGN12 can be driven by provision of acetate in laboratory culture, we next wanted to test if acetate could be used as the main carbon source in an applied setting. To this end, the performance of CGN12 and the non-ureolytic model MICP bacterium *A. pseudofirmus* were assessed in self-healing cement mortars using the previously established standard procedure (Tan et al. 2023). In brief, spores were produced for each strain and then encapsulated in lightweight aggregate as described in materials and methods. Encapsulated spores were cast into mortars with sodium acetate, calcium nitrate and trace amounts of yeast extract as nutrients dissolved in the mixing water, with concentrations corresponding to YAC medium composition. Mortars containing the same nutrients but no added spores (“Control”) and plain mortars containing neither nutrients nor bacteria (“Reference”) were produced for comparison. Once hardened, mortars were cured and then cracked under 3- point bending to obtain a target crack width of ∼0.5 mm. Following this, they were semi-submerged in tap water at room temperature and the self-healing process monitored over 8 weeks (Fig. 5A).

**Figure 5.**
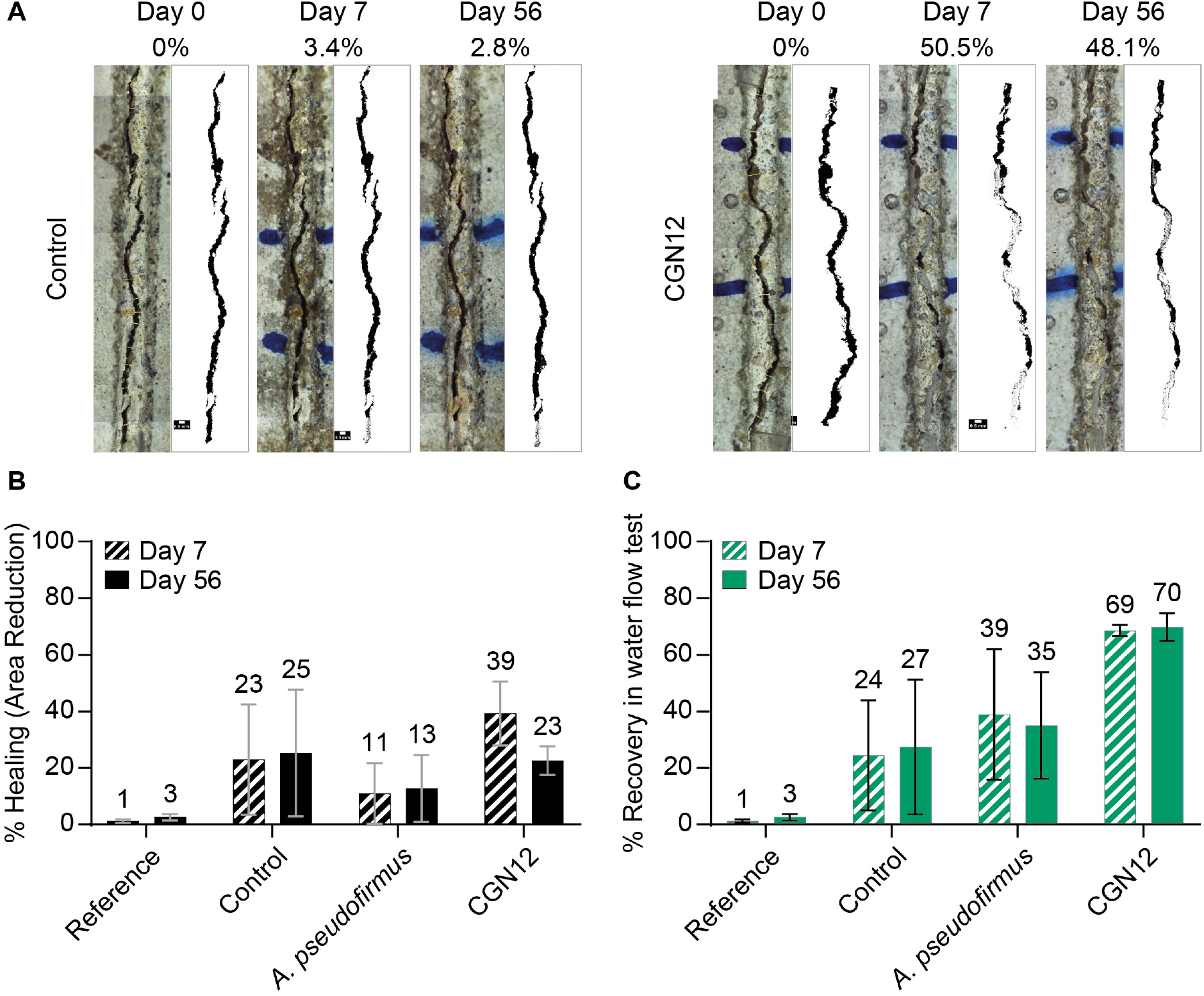
CGN12 efficiently seals cracks in mortar prisms. (A) Crack closure in control and CGN12- containing mortars with acetate provided as the main carbon source. Pictures were taken as a top-down view onto the surface of the specimen. A representative mortar sample is shown immediately after cracking and then after 7 and 56 days of healing. Digital rendering of the crack areas is shown to the right of each sample with the calculated percentage in crack area reduction shown above. Cracks were marked with blue pen to allow the same region to be monitored over time. Due to the limited imaging area of the microscope, images were manually assembled from multiple photographs of overlapping sections of the specimen. (B) Crack healing in mortars shown by average % area reduction ofcracks from digitally rendered images of duplicate samples. (C) Crack healing in mortars shown by average % recovery of watertightness determined by water flow tests in duplicate samples. Each dataset in panels B and C contains reference samples (no nutrients, calcium or spores), control samples (nutrients and calcium, but no spores), and samples containing nutrients, calcium and either A *pseudofirmus* or CGN12 spores.

Autogenous healing in the reference mortars containing standard components only was minimal with 1-3% crack area reduction over the 56 days period. Some self-healing (23-25%) was observed in the control mortars, where nutrients but no bacteria were added. This is a common observation in these experiments, as self-healing is not tested under sterile conditions and environmental bacteria are likely to facilitate low levels of healing using the nutrients and additional calcium ions provided (Tan et al. 2020). Mortars containing *A. pseudofirmus* spores showed a lower percentage of crack healing than the control mortars (13% after 56 days), consistent with the low MICP-activity of this strain when grown on acetate as the main carbon source. In contrast, CGN12 displayed greater crack healing than the control after 7 days (39%), although this subsequently reduced to 23% after 56 days (Fig. 5AB).

While quantification of the decrease in crack area between samples is a useful tool for monitoring the early stages of crack healing, it can only give visual information of the crack surface, but cannot account for healing product formed at the depth of the crack (He et al. 2023). The main aim of using biomineralization in self-healing concrete is to re-establish the water tightness of the structure, which is lost when microcracks form, in turn leading to corrosion of internal steel reinforcements and eventually structural failure. To assess self-healing performance, water flow assays can test the restoration of watertightness in cracked mortars, which can be mediated by healing product at both crack surface and deeper in the fissure. When we tested our mortars after 30 days and 56 days of healing, reference mortars (which contained no nutrients, calcium or spores) showed almost no recovery of watertightness of 1-3% (Fig. 5C), consistent with the crack area reduction data. Control mortars, which contained nutrients and calcium but no spores, showed up to 27% recovery of watertightness after 56 days, likely due to activity of environmental bacteria as mentioned above. While mortars containing *A. pseudofirmus* spores showed a high degree of variability between duplicates and only barely surpassed the control specimens, samples containing CGN12 achieved the greatest recovery of water tightness, averaging 69% after 30 days and 70% after 56 days (Fig. 5C).

Taken together, these results show that CGN12 can provide self-healing properties and restoration of water tightness when cast into cement mortars and provided with acetate as the main carbon source.

### The genetic basis for variation in calcite precipitation between species

The experimental evidence presented so far suggested that non-ureolytic MICP, at least in some bacteria, is driven by cellular metabolism of carbon sources such as acetate. To explore if the differences between species observed in the precipitation assays could be explained by differences in the metabolic enzymes each species possessed, a targeted genomic comparison was performed. For this, we primarily focussed on genes that were annotated as being involved in acetate metabolism. Additionally, we were interested in genes encoding putative calcium binding domains that could be exposed on the cell surface and might influence MICP, positively or negatively, via either creating a local calcium enrichment or sequestration of calcium ions.

To this end, we returned to the five strains from our initial set that showed a positive effect on MICP via increasing the acetate concentration: CGN12, BA32, PD1-1, UBN2 and Psy5 (Fig. 1). Genomic sequences for these strains were obtained through a combination of Illumina short read sequencing and Oxford Nanopore long-read sequencing. The high-fidelity short-read sequences were assembled using the long-read sequences as scaffolds, leading to nearly closed (2-12 contigs) draft genomes for these new isolates (Table S3). Published sequences for *A. pseudofirmus* and *S. cohnii* were obtained from NCBI (https://www.ncbi.nlm.nih.gov/) for comparison.

To analyse the gene content for acetate metabolism, we focused on genes encoding the enzymes of the tricarboxylic acid (TCA) cycle, as well as the enzymes needed to activate acetate for entry into the TCA cycle (Fig. 6A). For each of the seven species analysed, we identified presence or absence of the gene using the KEGG database and determined the copy number for each gene (Table 3, Table S4). In all analysed genomes, genes for the core metabolic pathway for acetate degradation via the TCA cycle were consistently present. All genomes included 2-3 copies of a gene encoding Acetyl-CoA-Synthase (ACS, EC 6.2.1.1), needed to feed acetate into the TCA cycle for degradation. Interestingly, however, the genome of CGN12 also contained a gene for an ADP-forming ACS (EC 6.2.1.13), which is normally only found in archaea. At the same time, its genome was the only one to lack genes for the enzymes phosphate transacetylase (PTA) and acetate kinase (ACK), encoding the enzymes used by most bacteria to produce acetate as a metabolic end-product. These differences may suggest the presence of a unique acetate metabolism in this strain.

**Figure 6.**
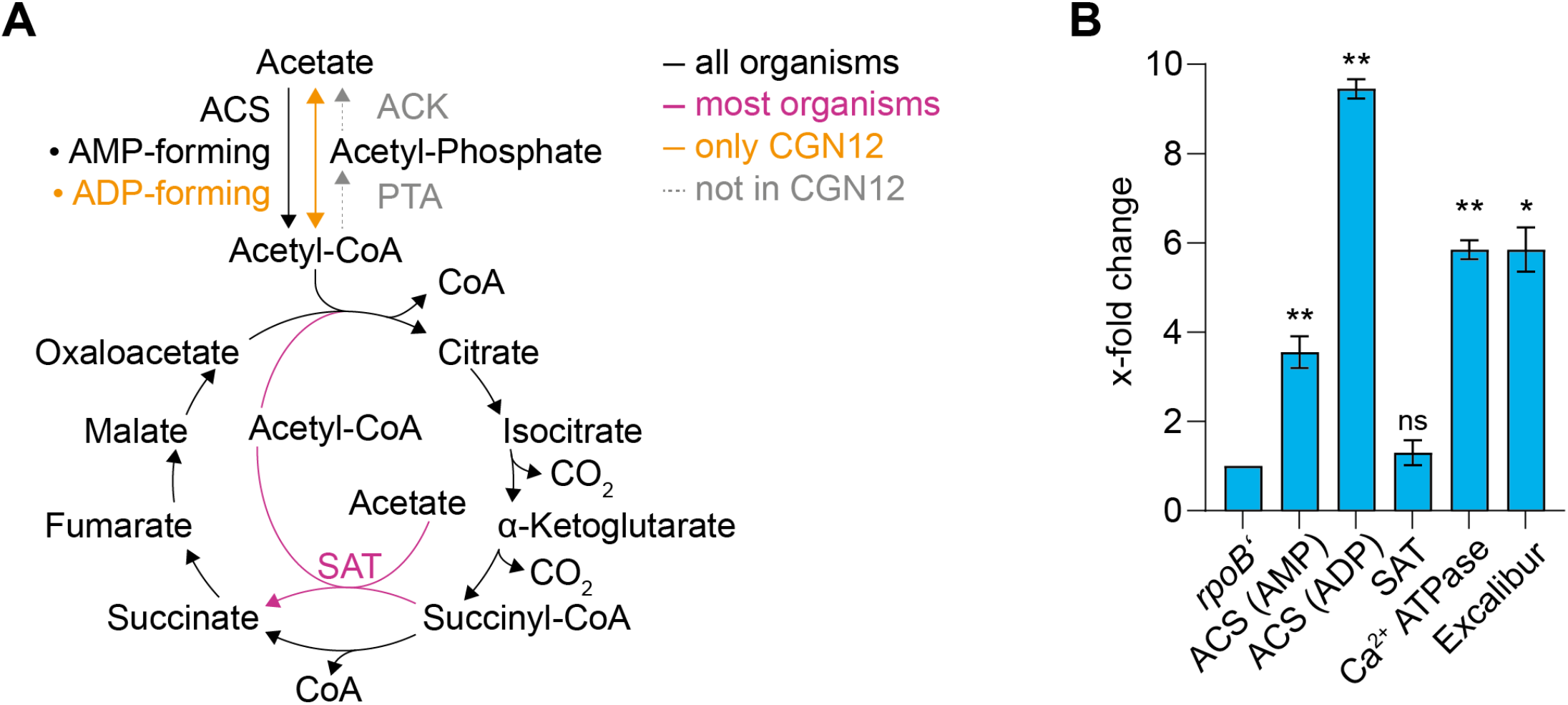
Genomic comparisons of isolated species and changes in gene expression in CGN12. A. Similarities and differences in how acetate can feed into the TCA cycle in selected species. ACS, Acetyl-CoA-Synthetase, showing two types of the enzyme, one forming AMP and the other forming ADP during the reaction; ACK, Acetate kinase; PTA, Phospho-transacetylase, SAT, Succinyl-CoA:Acetate CoA transferase. B. Changes of gene expression in CGN12 upon calcium challenge, determined by RT-qPCR. X-fold changes were calculated using the Pfaffl equation from biological duplicates and using the *rpoB*’gene as internal standard. Samples were taken before and 30 min after the addition of 100 mM calcium nitrate to stationary phase cells to mimic MICP conditions. *, P<0.01; **, P<0.0001; ns, not significant in two-sided Student’s t-test.

**Table 3.** Genomic comparison of the number of acetate and TCA cycle related genes in the selected species.^a^.

| Organism | Acetyl-CoA Synthetase (AMP-forming) | Acetyl-CoA Synthetase (ADP-forming) | Acetate kinase | Phosphate acetyltransferase | Succinyl-CoA:acetate CoA transferase |
| --- | --- | --- | --- | --- | --- |
| *A. pseudofirmus* | 3 | - | 1 | 1 | - |
| *S. cohnii* | 3 | - | 1 | 1 | 1 |
| BA32 | 2 | - | 1 | 1 | - |
| CGN12 | 2 | 1 | - | - | 2 |
| PD1_1 | 3 | - | 1 | 1 | - |
| Psy5 | 2 | - | 1 | 1 | 1 |
| UBN2 | 2 | - | 1 | 1 | 1 |

<sup>a</sup> For details of the locus tag for each identified gene, please refer to Table S4

Another observation in the acetate metabolic genes was the presence of a gene encoding a putative succinyl-CoA:acetate CoA transferase (SAT, EC 2.8.3.18) in several of the analysed species (Fig. 6A, Table 3, Table S4). This enzyme is involved in heterotrophic growth on acetate using a modified TCA cycle where conversion of succinyl-CoA to succinate is coupled to activation of acetate to acetyl-CoA in a single enzymatic reaction and with the expenditure of only a single ATP-equivalent (Pettinato et al. 2022). *S. cohnii*, Psy5 and UBN2 both possess one copy of such a gene, while CGN12 harbours two, highlighting yet another unique feature of its acetate metabolic genes.

In addition to acetate metabolism, we were also interested in the differences in calcium binding proteins between our strains. Tables 4 and S5 show the copy number and locus tags, respectively, of genes for selected calcium binding proteins: Excalibur domain proteins, which have a calcium binding motif but an unknown function (Rigden et al. 2003), as well as an F-type and a P-type calcium binding ATPase thought to be involved in calcium homeostasis through export (Norris et al. 1996). All analysed species possess a single copy of each gene encoding the two calcium ATPases. Genes coding for an Excalibur domain protein are present in BA32, CGN12, PD1_1 and UBN2, but such a gene is absent from *A. pseudofirmus*, *S. cohnii* and Psy5. Thus, in addition to differences in acetate metabolism, we also detected notable differences in calcium binding proteins that are likely to be surface exposed. Given that acetate metabolism provides DIC, and calcium binding proteins may affect the local concentration of calcium ions near the cell surface, it is conceivable that these genomic differences contribute to the differing MICP properties of the analysed strains.

**Table 4.** Genomic comparison of the number of selected calcium homeostasis related genes in the selected species.^a^.

| Organism | Excalibur domain proteins | F-type Ca<sup>2+</sup> ATPase | P-type Ca<sup>2+</sup> ATPase |
| --- | --- | --- | --- |
| *A. pseudofirmus* | - | 1 | 1 |
| *S. cohnii* | - | 1 | 1 |
| BA32 | 2 | 1 | 1 |
| CGN12 | 2 | 1 | 1 |
| PD1_1 | 1 | 1 | 1 |
| Psy5 | - | 1 | 1 |
| UBN2 | 1 | 1 | 1 |

### The effect of calcium on the gene expression of metabolic enzymes and calcium binding proteins

To better understand the role of the identified acetate metabolism and calcium binding domain encoding genes for MICP in CGN12, we determined the expression levels of some of these genes under MICP-conditions using quantitative Reverse Transcriptase PCR (qRT-PCR). As the earlier experiments had shown that acetate-driven MICP by CGN12 occurs mostly in stationary phase, we here created the relevant condition by exposing stationary-phase cells to a calcium stimulus (100 mM calcium nitrate) for 30 min. Untreated cells served as the control. The genes of interest were those encoding the two distinct types of ACS enzymes (gene locus AB1K09_14095 for EC 6.2.1.1, and locus AB1K09_11545 for EC 6.2.1.13), one of the genes encoding an SAT (gene locus AB1K09_18240), the P-type Calcium binding ATPase (gene locus AB1K09_00890), and one of the Excalibur domain proteins. For the latter, the gene with locus tag AB1K09_06620 was chosen, which contains only an Excalibur domain, whereas the second gene (AB1K09_02315) encodes a protein with an Excalibur, S-layer homology and metallo-beta-lactamase domains, which makes it more difficult to predict its potential function. As a constitutively expressed reference gene, the housekeeping gene *rpoB’* encoding the β’ subunit of RNA polymerase was used.

As expected, addition of calcium led to an increase in expression of the genes putatively involved in calcium export or binding, with the level of the P-type calcium ATPase gene increasing 5.9-fold and the Excalibur domain gene 5.8-fold compared to the untreated sample (Fig. 6B). The upregulation of the ATP-dependent calcium transporter upon calcium addition potentially reflects a need of the cell to remove excess calcium ions that passively entered the cytoplasm through active export. The role of the Excalibur domain protein is less clear and is discussed in more detail below.

Interestingly, both of the ACS-encoding genes showed an increase in expression, 3.5 and 9.5 times higher, respectively, in the treated condition compared to the untreated sample (Fig. 6B). The SAT, presumably involved in a modified TCA cycle, did not show any significant increase in expression in the presence of calcium and may therefore either play a minor or constitutive housekeeping role in acetate degradation. These findings suggest an increase in acetate catabolic activity in response to calcium, which may be required to meet the increased energy demand from the upregulation of the ATPase. Such an up-regulation of acetate degradation could potentially explain the slightly higher degree of acetate consumption we had observed for CGN12 when grown in high-calcium conditions (Fig. 2B).

### Deletion of the putative calcium-binding Excalibur domain protein in *S. silvestris* CGN12

Our RT-qPCR data showed a clear upregulation of a gene encoding a putative calcium-binding Excalibur domain protein upon a calcium stimulus. The function of this protein or indeed the Excalibur domain in general is not yet known. To further investigate this gene’s putative role in MICP or calcium homeostasis, we generated an unmarked deletion of the excalibur gene (Δ*exc*) in CGN12 via double homologous recombination. To this end, the well-established vector pMAD, possessing a temperature-sensitive origin of replication and used for construction of unmarked gene deletions in *B. subtilis* (Arnaud et al. 2004), was modified with an origin of transfer (*oriT*) from plasmid pG2k-orit-gfp. The resulting plasmid, pMAD-oriT, could replicate in CGN12, and the *oriT* addition facilitated conjugation from *E. coli* donor strains to CGN12. Despite the lower routine growth temperature of CGN12 (30°C) compared to *B. subtilis* (37°C), the temperature-shifts required to control vector replication and thus the two homologous recombination events leading to gene deletion could be applied with minor modifications of the established protocols as described in the methods section.

Once the Δ*exc* strain had been obtained, we compared its calcium carbonate precipitation to that of the parental wild type. Overall, we did not observe a marked alteration in precipitation yields between the two strains at the previously used calcium and acetate concentrations (Fig. 7). We did observe a small but reproducible shift in the CaCO3-to-acetate stoichiometry from 0.47±0.01 in WT to 0.54±0.01 in the Δ*exc* strain, indicating that the latter might form slightly more calcium carbonate per acetate consumed than the wild type. Given how consistent the stoichiometry was for CGN12 between repeat experiments and different growth conditions, this 15% shift may be biologically meaningful and imply a role of the Excalibur-domain protein in calcium binding, potentially sequestering the ions and making them unavailable for crystal formation. Further investigations of this protein will be required to determine if it really contributes, positively or negatively, to non-ureolytic MICP in CGN12, or other bacteria with such genes.

**Figure 7.**
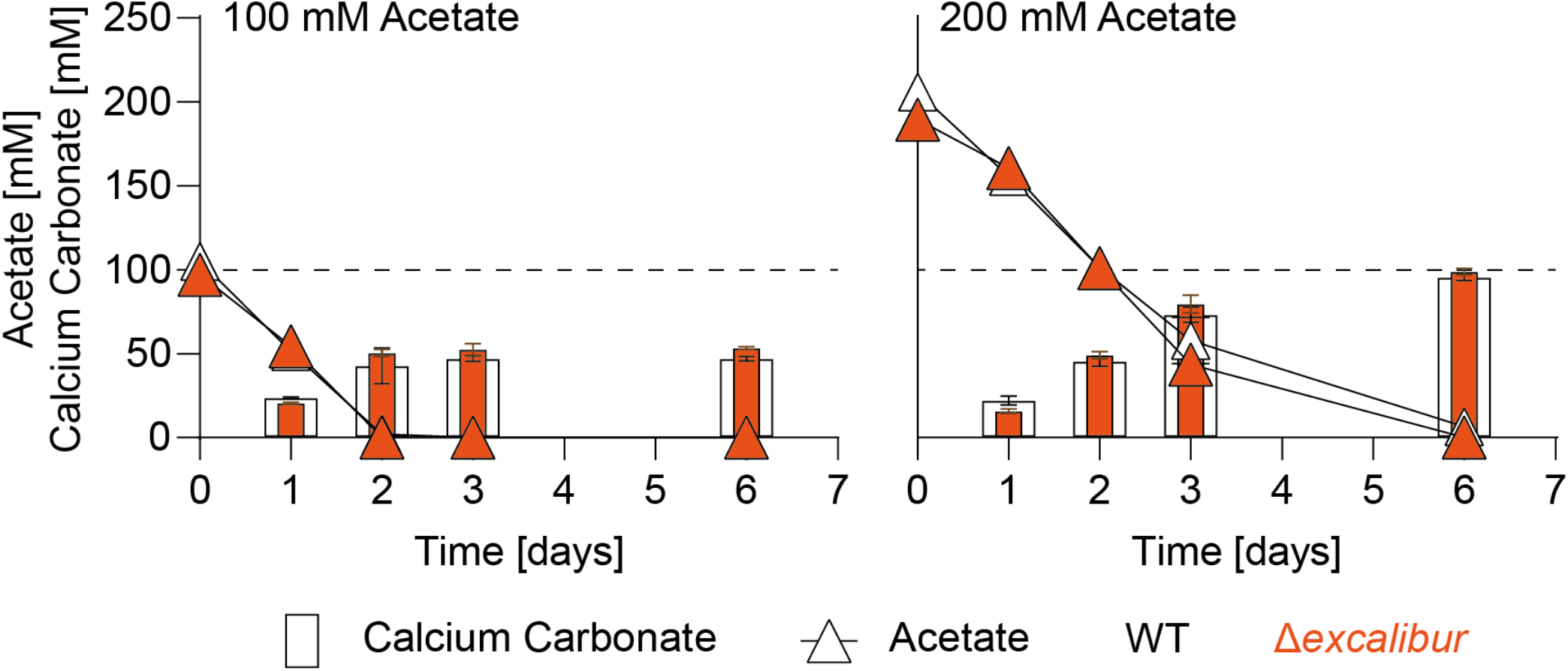
Cells lacking the Excalibur domain protein are not affected in MICP or acetate metabolism. Calcium carbonate precipitation and acetate consumption were determined for wild-type CGN12 and the Δexc deletion strain grown at different concentrations of sodium acetate. The initial concentration of calcium nitrate was 100mM in standard YAC medium. Cells were grown over 6 days at 30°C with shaking (140rpm), and one entire flask culture was sacrificed for each time course for quantification of precipitate formed and acetate remaining. Dashed horizonal lines indicate the theoretical maximum calcium carbonate precipitation. Data are show as mean and standard deviation of the mean for two biological replicates.

## Discussion

MICP serves as an exciting approach for more sustainable infrastructure materials. While ureolytic MICP is well-understood, there is only little knowledge on the non-ureolytic pathway(s) leading to biomineralization. This study aimed to give insights into the metabolic processes of non-ureolytic MICP by investigating several environmental isolates. We could show that in certain species, provision of acetate as a carbon source can be a major driver of mineral formation. Additionally, full genome sequencing, gene expression analysis and development of basic molecular techniques was used to shed some light on the underpinning processes and genetic features that contribute to non-ureolytic MICP.

Specifically, in the isolate *S. silvestris* CGN12, acetate consumption was directly linked to MICP, even though only a small proportion of the provided acetate was used by the bacteria for cell growth. In fact, the majority of acetate consumption occurred during the stationary growth phase, which was also the phase in which detectable MICP took place. Moreover, we observed a consistent carbonate-to-acetate ratio of 0.47-0.48 in the precipitation assays, suggesting that a quarter of the acetate-derived carbon was bound into CaCO₃, allowing *S. silvestris* CGN12 to precipitate almost all of the supplied calcium, as long as enough acetate was available. Our findings are in accordance with previous reports that suggested that non-ureolytic MICP is governed by DIC and the available calcium ions, as well as suitable pH values and presence of nucleation sites (Hammes and Verstraete 2002; Hoffmann et al. 2021). Together, these observations indicate that non-ureolytic MICP can occur independently of active cell growth, with acetate catabolism leading to DIC formation to fuel the MICP process.

Comparing the different environmental isolates and laboratory strains showed that the metabolic link between acetate consumption and MICP was not universal among all non-ureolytic MICP bacteria. The observed variations suggest that different species may utilize alternative metabolic pathways or carbon sources to drive DIC production, leading to MICP. Understanding these metabolic nuances will be crucial for the application of MICP technologies in the future, because it allows a precise design of nutrient provision to maximise mineral formation by a given bacterial strain.

When testing *S. silvestris* CGN12 in an application-relevant context, our data showed that acetate can serve as an effective nutrient for crack healing in cement mortars. This is a significant step towards reducing the cost of self-healing concrete using non-ureolytic bacteria via replacing the majority of expensive yeast extract with the much cheaper sodium acetate. Use of acetate also avoids the drawbacks associated with more complex media, such as retarding effects on cement hydration and potential adverse impacts on the final material (Schreiberová et al. 2019). The sodium salt of acetate is cost-effective and has been demonstrated to enhance the rigidity of concrete by improving interfacial bonds between hydration products and aggregates (Al-Kheetan et al. 2020). Our results further show that physiological experiments in the laboratory can provide useful insights to optimise the application conditions for MICP-based technologies.

Genomic comparison between our isolates to better understand the differences in acetate metabolism and its link to MICP, revealed an intriguing difference in the pathway feeding acetate into the TCA cycle in the genome of CGN12. In bacteria, acetate activation to acetyl-CoA usually occurs via the enzyme Acetyl-CoA-Synthase (EC 6.2.1.1), which uses the energy from ATP hydrolysis to AMP and pyrophosphate to produce acetyl-CoA. This reaction is considered irreversible under physiological conditions, and bacteria producing acetate as a metabolic end-product or during overflow metabolism generally employ the two sequential reactions of phosphate transacetylase (PTA), converting acetyl-CoA to acetyl phosphate, and then acetate kinase (ACK), converting acetyl-phosphate to acetate, with concomitant generation of ATP. Curiously, CGN12 does not possess genes encoding these two enzymes, but instead harbours a gene encoding an ADP-forming ACS (EC 6.2.1.13). This enzyme is almost exclusively found in archaea such as *Thermococcus* and in some protists, but only seldomly in bacteria (Schönheit and Schäfer 1995; Reeves et al. 1977; Parizzi et al. 2012; Awano et al. 2014). It catalyses the reversible conversion between acetate and acetyl-CoA and can therefore be used both in acetate production and consumption, although in Archaea it is mostly linked to the former (Schönheit and Schäfer 1995). The strong induction of the ADP-forming enzyme we observed in CGN12 under calcium exposure suggests that its role under these conditions is likely catabolic, i.e. activation of acetate to acetyl-CoA, in contrast to reports from other bacteria that may be using the enzyme predominantly in the reverse direction (Parizzi et al. 2012). The unusual presence of two different ACS, but no ACK or PTA in CGN12, as well as the presence of two succinyl-CoA:acetate CoA transferase (SAT) encoding genes, may indicate that acetate metabolism generally plays a unique role in the physiology of this bacterium, possibly beginning to shed some light on its efficient MICP ability using acetate as the carbon source.

Intriguingly, in *S. silvestris* CGN12 exposure to high calcium levels lead to the upregulation not only of genes related to calcium detoxification, but also of those involved in acetate metabolism. Together with our observation that under high calcium conditions, CGN12 appeared to consume slightly more acetate, this may indicate a direct link between calcium stress and acetate metabolism and may suggest that MICP serves as an active calcium detoxification mechanism in this species. Such a mechanism might be biologically plausible, given the natural habitat of the bacterium, i.e. calcium-rich limestone rock, from which it was originally isolated (Reeksting et al. 2020).

Further support for this notion may be found in previous studies in micro-algae, where it was found that extracellular precipitated calcium could originate from within a microorganism, leading the authors to propose that carbonatogenesis was an active process involving ionic exchanges through the cell membrane (McConnaughey and Whelan 1997; Castanier et al. 1999; Yates and Robbins 1999). Consistent with this, we could show that a gene encoding a calcium-binding P-type ATPase in *S. silvestris* CGN12 was significantly upregulated upon calcium exposure. These findings support a model in which excess intracellular calcium is actively exported from the cell and, using CO2 from acetate metabolism, is incorporated into extracellular calcium carbonate on the cell surface (Fig. 8), consistent with the hypothesis formulated by Hammes and Verstraete (Hammes and Verstraete 2002). In how far extracellular calcium binding sites are involved in MICP, and whether they promote or hinder biomineralization remains to be fully explored. CGN12 possesses two proteins with a predicted Excalibur calcium binding domain. Our original hypothesis had been that presentation of calcium ions on the cell surface by such proteins might aid in crystal nucleation. However, the MICP-yield of a strain lacking one of these proteins was almost identical to the parent strain, with the main difference being a small but reproducible increase in the carbonate-to-acetate ratio, which may indicate that the protein may rather sequester the calcium and lower the MICP efficiency. More detailed mechanistic studies will be needed to answer these questions.

**Figure 8.**
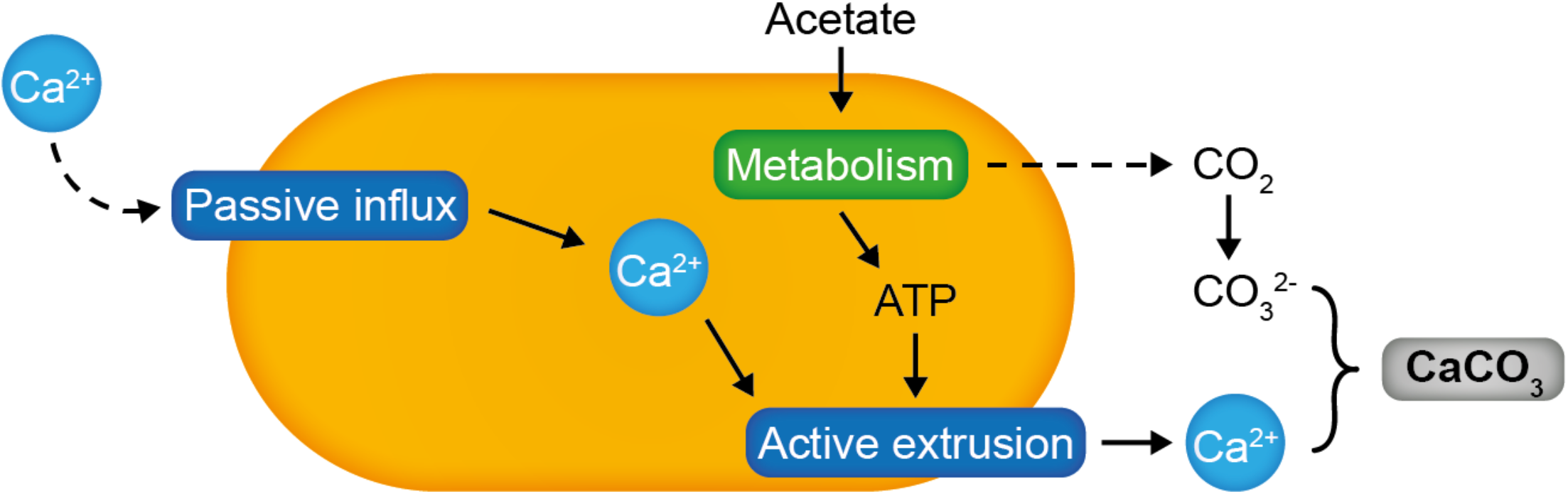
Model of non-ureolytic MICP in *S. silvestris* CGN12. Calcium enters the cell passively and needs to be actively exported. The energy for this process is provided by acetate catabolism, which also produces CO2 as a byproduct. The CO2 in solution can react with calcium ions to form CaCO3, using the bacterial cell surface as a nucelation site.

To summarize, the direct link between carbon metabolism and MICP in *S. silvestris* CGN12 reported here provides valuable insights into the physiological and molecular mechanisms of non-ureolytic biocalcification. While not a universal principle across all non-ureolytic MICP strains, knowledge of this relationship offers a means to control MICP through acetate provision, making it a powerful tool in biotechnological applications. Moreover, we provide initial indications of potential genetic determinants of MICP activity, paving the way towards gaining a full understanding of non-ureolytic biomineralization.

## Material & Methods

### Strains and growth conditions

All bacterial strains and plasmids used in this study are listed in Table 1 and S1. Environmental strains were previously isolated from limestone rock scrapings (Reeksting et al. 2020). The psychrotrophic strain Psy5 was isolated as described but incubated at 7.5°C. Unless otherwise stated, bacteria were grown in Yeast extract Acetate (YA) medium containing 50 mM Tris-HCl pH 7.8, 0.2% w/v yeast extract, 0.5mM MgSO4, 0.01mM MnSO4 and 100 mM sodium acetate. YA medium containing 200 mM sodium acetate was used when indicated. Where indicated, cells were grown in LB pH 8.2, adjusted with a final concentration of 20 mM M Tris-HCl pH 9.

*E. coli* strains were routinely grown in regular LB at 37°C with shaking (140 rpm).

### Endospore production

Endospores of CGN12 and *A. pseudofirmus* were prepared in Difco sporulation medium (DSM; 8 g · liter^-1^ nutrient broth, 13.41 mM KCl, and 0.49 mM MgSO4, adjusted to pH 7.6 with NaOH. Prior to use, 1 mM Ca(NO3)2, 0.01 mM MnCl2, and 1µM FeSO4 were added from a filter-sterilized stock solution (Sonenshein et al. 1974). Cultures were grown in LB for 24 h at 30°C with shaking (140 rpm) were used to inoculate 750 ml of sporulation medium to an OD600 of 0.02. Cultures were grown for at least 48 h before sporulation was assessed using phase-contrast microscopy. When most cells contained phase-bright endospores, cells were pelleted by centrifugation (4,000xg, 10 min at RT) and washed three times in 10 mM Tris-HCl pH 9. Cells were treated with chlorhexidine digluconate (0.3mg ml^-1^) for 30 min to kill vegetative cells, and then the wash steps repeated. Spore pellets were frozen at -80°C and freeze-dried under vacuum overnight.

### Precipitation assays

Cultures grown for 24 h in YA medium were used to inoculate 20 ml YAC medium, containing 50 mM – 100 mM calcium nitrate, to an OD600 of 0.02. Precipitation cultures were incubated with shaking (140 rpm) at 30°C. At the indicated timepoints, precipitate was harvested by centrifugation (500 × *g*, 5 min at RT). The supernatant containing the cells was discarded and the precipitate washed in distilled water until all remaining cells were removed. The precipitate was dried at 50°C for 3 days until the weight remained constant. The amount of precipitate was determined as the difference in weight of each empty tube before harvesting and the post-drying weight. Separate flasks per time point were set up so that at each time point (day 0, 1, 2,3 and 6 post inoculation) the calcium carbonate from a complete flask could be harvested and quantified.

### Determination of acetate concentration by HPLC

HPLC was performed on an Agilent 1260 Infinity at 65°C with a Rezex ROA Organic Acid H^+^ 8% column (Phenomenex). The mobile phase was 5 mM H2SO4, with a flow rate of 0.6 ml/min over 25 min. The primary detection wavelength was 215 nm, and the UV spectrum of each sample was collected. Samples were passed through a 0.22 µm filter and diluted 1:10 before 10 µL was injected onto the column. Acetate for each sample was quantified by comparing against a standard curve of samples with known acetate concentrations.

### Determination of soluble calcium concentration

Samples were centrifuged at 4,000 × *g* for 10 min to remove cells. The supernatant was passed through a 0.22 µm filter before 0.5 ml was combined with 2 ml of Patton-Reeders solution (0.105 mM calconcarboxylic acid, VWR, in 0.73 M NaOH) and vortexed briefly to mix. The sample was titrated with 25 mM EDTA until the solution turned a permanent bright blue, indicating that all calcium had been complexed. The volume of EDTA required to reach the colour change was recorded and used to calculate the concentration of calcium in the sample, based on EDTA complexing calcium in a 1:1 molar ratio.

### Whole genome sequencing

To extract genomic DNA, strains were grown for 24 h in LB pH 8.2. Cells were pelleted and genomic DNA extracted using the Monarch Genomic DNA Purification Kit (New England Biolabs). To obtain high molecular weight gDNA, after incubation with RNase A, sodium acetate was added to a final concentration of 300 mM and the entire solution was mixed with equal parts isopropanol. The precipitated DNA was then spooled on a glass rod and transferred into 100 µl Tissue Lysis Buffer mixed with 113 µl 10 mM Tris-HCl pH 8, before continuing with DNA binding to the column and elution as described by the manufacturer.

Short read genomic sequences were obtained by Illumina sequencing (Microbial Genome Sequencing Centre, Pittsburg, PA, USA). Paired end reads were assembled into contigs using Shovill (https://github.com/tseemann/shovill). Genome annotation was performed using the NCBI Prokaryotic Genome Annotation Pipeline (https://www.ncbi.nlm.nih.gov/genome/annotation_prok/).

Sequencing libraries for long read sequencing of the strains CGN12, BA32, PD1-1, UBN2 and Psy5 were generated from 400 ng high molecular weight gDNA per strain using the Oxford Nanopore Technologies Native Barcoding Kit 24 (SQK-NBD112.24) according to the manufacturers’ protocol. Quality control was performed using Invitrogen^TM^ Qubit^TM^ 4 Fluorometer. Next, 11 fmol of sequencing library with an estimated fragment size of 7.5 kb were loaded onto a R10.4 MinION flow cell (FLO-MIN112) and sequenced on a GridION X5 Mk1 for 72 h using the ‘barcode balancing’ option. Subsequent base calling was performed using Guppy v6.3.9 with super high accuracy (SUP) mode resulting in 125 - 233 k reads per strain.

Genome assembly was performed two-fold, with Unicycler v0.5.0 (Wick et al. 2017) for short- and long-read hybrid assembly and epi2me wf-bacterial-genomes v0.4.0 (Pedersen and Quinlan 2018; Kolmogorov et al. 2019) for long-reads only. Quality and completeness of assemblies was evaluated using common assembly metrics (number and size of scaffolds, N50, N90) and BUSCO v5.0.0 in genome mode with Bacillales database (Manni et al. 2021) and the best assembly for each strain selected. Gene annotation was performed using Prokka v1.14.6 (Seemann 2014). Completeness of the annotation was performed by extracting all protein sequences of annotated genes and comparing them with BUSCO in protein mode.

### RT-qPCR

Stationary-phase cells were treated either with or without 100 mM calcium nitrate for 30 min. RNA was isolated using a Total RNA Extraction kit (NEB #T2010). For reverse transcription, 2 µg of the isolated RNA was used to synthesize cDNA from random primers using a High Capacity cDNA RT Kit (Applied Biosystems). LuminoCt SYBR Green qPCR Readymix (Sigma Aldrich) was used for all qPCRs. Each reaction contained 10 μl of Master Mix, 0.2 μl of 100x ROX internal reference dye, 0.3 μM of each primer, 4 μl of diluted cDNA template and ultrapure water to a final volume of 20 μl. qPCR was performed on StepOnePlus^TM^ Instrument (Applied Biosystems). The qPCR cycle consisted of an initial enzyme activation step at 95°C for 20 s, followed by 40 cycles of denaturation at 95°C for 3 s and annealing and extension at 60°C for 30 s. A melt curve analysis was performed by raising the temperature at the end of each run by 0.3°C from 60°C to 95°C. Standard curves were created for each primer set with qPCR, using a dilution series of pooled cDNA from individually synthesized samples as a template. Fold changes were calculated using the Pfaffl equation (Pfaffl 2001).

### Deletion of the Excalibur protein-encoding gene in *S. silvestris* CGN12

The temperature-sensitive *oriT* was amplified from pG2k-orit-gfp using primers SG1073 and SG1073 (Table S2) and cloned into pMAD using the BamHI restriction sites (Arnaud et al. 2004). Genomic DNA was extracted from CGN12 (Genejet genomic DNA extraction Kit, Thermo, USA). Flanking regions 1 kB upstream and downstream of the Excalibur domain-encoding gene (gene locus AB1K09_06620) were amplified using primers SG1229 and SG1230 (upstream) and SG1231 and SG1232 (downstream). Fragments were joined by Gibson Assembly (NEBuilder) and ligated into pMAD-OriT using the NcoI restrictions sites. For the conjugation to CGN12, *E. coli* S17 containing the plasmid were grown into stationary phase in LB medium containing 100ug/ml ampicillin. CGN12 was also grown into stationary phase in LB pH 8.2 medium. S17 and CGN12 cells were washed and combined in a 1:1 ratio in 20 ml LB supplemented with 20 mM MgCl2. Cells were incubated for 1 h at 30°C with shaking and then pelleted by centrifugation and spotted onto LB agar plates containing 10 mM MgCl2. Plates were incubated at 30°C for 24 h. Cells were scraped off the agar plates, resuspended in 1 ml LB pH 8.2 and serially diluted in 1:10 steps. Dilutions were spotted in a volume of 20 µl per spot onto LB pH 8.2 agar containing MLS selection (0.5 µg/ml Erythromycin and 12.5 µg/ml Lincomycin). Plates were incubated for 72 h at 30°C and colonies patched onto fresh selection plates to confirm growth. Presence of the plasmid in CGN12 was confirmed by colony PCR with primers SG1072 and SG1073. Next, an overnight culture was prepared in LB medium with MLS selection at 30°C. The following morning, 10 ml LB with MLS selection were inoculated to a starting OD600 of 0.1 and incubated at 30°C for 2 h. The temperature was then increased to 42°C for 5 h. Serial dilutions were then plated onto LB agar with MLS selection and incubated at 42°C for 24 h. Colonies were selected and inoculated into in LB pH 8.2 without antibiotics and incubated for 6 h at 30°C. The temperature was then increased to 42°C for 3 h. Dilutions were plates onto LB pH 8.2 agar without selection and incubated for 24 h at 42°C. Colonies were replica-patched onto MLS and non-MLS plates. MLS-sensitive colonies were checked for presence of the gene deletion by PCR using primers SG1229 and1232. Positive clones were confirmed by sequencing of the PCR-product with primer SG1409.

### Strain and genome sequence deposition

The five strains chosen for full genome sequencing were deposited into the Deutsche Sammlung von Mikroorganismen und Zellkulturen (DSMZ) collection (Germany) under the following accession numbers: DSM 118343 (*M. dongyingensis* BA32), DSM 118343 (*S. silvestris* CGN12), DSM 118343 (*P. simplex* Psy5), and DSM 118343 (*P. frigoritolerans* UBN2). *B. licheniformis* PD1_1 was already deposited as DSM 110495 (Reeksting et al. 2020).

The hybrid genome sequences were deposited into GenBank with the following accession numbers: JBFEAO000000000 (*M. dongyingensis* BA32), JBFEAN000000000 (*S. silvestris* CGN12), JBFEAM000000000 (*B. licheniformis* PD1_1), JBFEAL000000000 (*P. simplex* Psy5), and JBFEAK000000000 (P. *frigoritolerans* UBN2)

### Preparation of mortar samples

Mortar prisms were prepared according to (Reeksting et al. 2020). Mortar prisms (65 mm by 40 mm by 40 mm) were cast in duplicate, in two separate layers. The first layer contained (per 2 prisms), 177.8 g sand conforming to standard BS EN 196-1, 61.3 g Portland limestone cement (CEM II/A-L 32.5R), 30.7 g water, 0.7 g yeast extract, 3.0 g calcium nitrate, and 2.4 g aerated concrete granules. Spores (1.0 x 10^10^ CFU) were resuspended in 2.83 g distilled water and soaked into the aerated concrete granules before drying and sealing with polyvinyl acetate (30% w/w). After 3 h, the second, top layer was cast. The second layer contained standard cement mortar (per 2 prisms: 184 g standard sand, 61.32 g cement, and 30.7 g water). Reference specimens were cast in two layers but contained only standard cement mortar in both layers. Specimens remained at room temperature for 24 to 48 h before demoulding and subsequent curing for 28 days submersed in tap water. After curing, specimens were oven dried at 50°C for 24 h. The top third of the prism was wrapped with carbon fibre-reinforced polymer strips to enable generation of a crack of controlled width. A notch (1.5 mm deep) was sawn at midspan to serve as a crack initiation point. Specimens were cracked by 3-point bending using a 30-kN Instron static testing frame. A crack mouth opening displacement (CMOD) gauge was used to measure crack width. Load was applied to maintain a crack growth of 25 µm per minute, and loading was stopped when the crack width was predicted to be 500 µm wide after load removal. A marker pen was used to indicate specific crack sections to enable monitoring of the crack at the same site. Following cracking, prisms were placed in tanks that were open to the atmosphere and filled with tap water to 20 mm below the top of the mortars and then incubated at room temperature for 2 months. Visualization of crack healing was monitored using a Leica M205C light microscope, and images were taken of freshly cracked mortars and after 1, 4, and 8 weeks of healing.

## Acknowledgements

This work was supported in part by funding from the Engineering and Physical Sciences Research Council (EPSRC, UK) through the Resilient Materials for Life (RM4L) (EP/P02081X/1) project and the Engineering Microbial-Induced Carbonate Precipitation via Meso-Scale Simulations (eMICP) (EP/S013997/1; EP/S013997) project.

The authors gratefully acknowledge the Technical Staff within the Life Sciences Department and the Department of Architecture and Civil Engineering at the University of Bath for technical support and assistance in this work. We also thank Yannic Zettelmeyer at Johannes-Gutenberg University Mainz for genome assembly from the long and short read data as part of his student research project.

## Data accessibility statement

The data that support the findings of this study are available in the supplementary material.

