## Supplemental tables for "Metabolic insights into microbially induced calcite formation by Bacillaceae for application in bio-based construction materials"

**Supplementary data**

**Table S1**. Strains, vectors and plasmids used in this study

**Table S2**. Oligonucleotides used in this study

**Table S3**. Overview of whole genome sequencing data of five environmental isolates

**Table S4**. Genomic comparison of acetate and TCA cycle related genes

**Table S5**. Genomic comparison of selected calcium homeostasis genes

**Table S6**. Experimental numerical data for Figures 1-8, available as separate Excel file

**References**

**Table S1**. Strains, vectors and plasmids used in this study. The used environmental isolates are listed in Table 1.

| **Strains** | **Description** | **Source** |
| --- | --- | --- |
| *A. pseudofirmus* | WT | (Nielsen et al. 1995) |
| *S. cohnii* | WT | (Spanka and Fritze 1993) |
| *E. coli* DH5α | *supE44 ΔlacU169(Φ80lacZΔM15)*  *hsdR17 recA1 endA1 gyrA96 thi-1 relA1* | Laboratory stock |
| *E. coli* S-17 | *TpR SmR recA thi pro*  *hsdR-M+RP4: 2-Tc:Mu:Km Tn7 λpir* | Laboratory stock |
| **Vectors** |  |  |
| pMAD | for generating unmarked deletion/insertion mutants in Gram+ bacteria | (Arnaud et al. 2004) |
| **Plasmids** |  |  |
| pG2k-oriT-gfp | contains origin of transfer | David Leak, University of Bath |
| pMAD-oriT | pMAD containing the origin of transfer from pG2K-oriT-gfp | This study |
| pMADOriTExcalibur | pMAD-oriT containing the deletion construct for Δ*excalibur* | This study |

**Table S2**. Oligonucleotides used in this study

| **Name** | **Description** | **Sequence 5’ → 3’ *^a^*** |
| --- | --- | --- |
| SG1072 | oriT cassette + BamHI fwd | TTTAA**GGATCC**TCTTCTTGATGGAGCGCATG |
| SG1073 | oriT cassette + BamHI rev | TTTAA**GGATCC**CGCACGATATACAGGATTTTG |
| SG1229 | Up-fwd Excalibur NcoI | AATTT**CCATGG**GCAGATGAAACATTC |
| SG1230 | Up-rev Excalibur | ***ACAACTCCTA***CATTGGAAAAAACCTCCCAAAC |
| SG1231 | Do-fwd Excalibur | ***TTTTCCAATG***TAGGAGTTGTGAAATAGATGAAAATTCTC |
| SG1232 | Do-rev Excalibur NcoI | AATTT**CCATGG**GCTGAGCAATCATTTC |
| SG1253 | qPCR ACS1 (EC 6.2.1.13) fwd | CAGCCCAGTATGGCATCAA |
| SG1254 | qPCR ACS1 (EC 6.2.1.13) rev | AGCCGCTGATCACTTCAATC |
| SG1257 | qPCR Excalibur fwd | TGGAAGTAGAAGGAGCTGCAAA |
| SG1258 | qPCR Excalibur rev | TTCCGCCTTTGTTCGTTGT |
| SG1259 | qPCR SAT fwd | AAATGCTGAGCAACGTATTGG |
| SG1260 | qPCR SAT rev | ATTAATGAAGGCGCATCTGG |
| SG1261 | qPCR ACS2 (EC 6.2.1.1) fwd | CGCAGTTCACAAATCGACAC |
| SG1262 | qPCR ACS2 (EC 6.2.1.1) rev | CATTACGGCCTTCAAACCAG |
| SG1279 | qPCR RpoB' fwd | ACGATAACCGAACGACCAGA |
| SG1280 | qPCR RpoB' rev | GGTCCTGGTAACCGTCCTTT |
| SG1301 | qPCR Calcium ATPase fwd | AGGACGAGCAAGGCTTAACA |
| SG1302 | qPCR Calcium ATPase rev | CAGGTTTAATTCCCGCTTCA |
| SG1409 | Excalibur knockout check F | GGACATTGGCAGCAAAAGG |

*^a^* Restriction sites are in uppercase bold; overhangs for the Gibson Assembly are in uppercase bold and italicised

**Table S3**. Overview of whole genome sequencing data of five environmental isolates.

|  | **BA32** | **CGN12** | **PD1_1** | **Psy5** | **UBN2** |
| --- | --- | --- | --- | --- | --- |
| Total Mb | 5.73 Mb | 4.04 Mb | 4.56 Mb | 5.77 Mb | 6.03 Mb |
| GC content % | 39.7% | 38.7% | 45.7% | 39.9% | 40.3 |
| Contigs | 12 | 2 | 2 | 2 | 3 |
| Mean Contig Coverage | 52.5x | 134x | 68x | 44x | 53x |
| Contig size | 5.12 Mb, 474 Kb, 76 Kb, 58 Kb, 2 Kb, 1 Kb, 1 Kb, 0.5 Kb, 0.5 Kb, 0.4 Kb, 0.4 Kb, 0.3 Kb | 3.97 Mb, 71 Kb | 4.49 Mb, 72 Kb | 5.53 Mb, 239 Kb | 5.77 Mb, 242 Kb, 10 Kb |
| Genbank Accession | JBFEAO000000000 | JBFEAN000000000 | JBFEAM000000000 | JBFEAL000000000 | JBFEAK000000000 |

**Table S4**. Genomic comparison of acetate and TCA cycle related genes, given as locus tag identifiers, in selected species.

| **Organism** | **Acetyl-CoA Synthetase (AMP-forming)** | **Acetyl-CoA Synthetase (ADP-forming)** | **Acetate kinase** | **Phosphate acetyltransferase** | **Succinyl-CoA:acetate CoA transferase** |
| --- | --- | --- | --- | --- | --- |
| *A. pseudofirmus* | WEG16239; WEG16252; WEG17604 | - | WEG16230 | WEG16846 | - |
| *S. cohnii* | WP_066413159; WP_066420872; WP_094366011 | - | WP_066421356 | WP_066421944 | WP_066418176 |
| BA32 | AB1K32_00085; AB1K32_27600 | - | AB1K32_00130 | AB1K32_21205 | - |
| CGN12 | AB1K09_14095; AB1K09_14185 | AB1K09_11545 | - | - | AB1K09_09305; AB1K09_18240 |
| PD1_1 | AB1K19_04130; AB1K19_04215; AB1K19_10845 | - | AB1K19_04270 | AB1K19_24055 | - |
| Psy5 | AB1K18_20090; AB1K18_20145 | - | AB1K18_20000 | AB1K18_26695 | AB1K18_02865 |
| UBN2 | AB1K12_20115; AB1K12_201780 | - | AB1K12_20030 | AB1K12_27310 | AB1K12_02960 |

**Table S5**. Genomic comparison of selected calcium homeostasis genes, given as locus tag identifiers, in selected species.

| **Organism** | **Excalibur domain proteins** | **F-type Ca^2+^ ATPase** | **P-type Ca^2+^ ATPase** |
| --- | --- | --- | --- |
| *A. pseudofirmus* | - | WEG16779 | WEG15677 |
| *S. cohnii* | - | WP_066418346 | WP_066421864 |
| BA32 | AB1K32_18280; AB1K32_22100 | AB1K32_21550 | AB1K32_09520 |
| CGN12 | AB1K09_06620; AB1K09_02315 | AB1K09_03070 | AB1K09_00890 |
| PD1_1 | AB1K19_23255 | AB1K19_00170 | AB1K19_12595 |
| Psy5 | - | AB1K18_26420 | AB1K18_08120 |
| UBN2 | AB1K12_21980 | AB1K12_27045 | AB1K12_08175 |

**Table S6**. Experimental numerical data for Figures 1-8, available as separate Excel file
